# Powerful and Efficient Strategies for Genetic Association Testing of Symptom and Questionnaire Data in Psychiatric Genetic Studies

**DOI:** 10.1101/383471

**Authors:** Aaron M. Holleman, K. Alaine Broadaway, Richard Duncan, Lynn M. Almli, Bekh Bradley, Kerry J. Ressler, Debashis Ghosh, Jennifer G. Mulle, Michael P. Epstein

## Abstract

Genetic studies of psychiatric disorders often deal with phenotypes that are not directly measurable. Instead, researchers rely on multivariate symptom data from questionnaires and surveys like the PTSD Symptom Scale (PSS) and Beck Depression Inventory (BDI) to indirectly assess a latent phenotype of interest. Researchers subsequently collapse such multivariate questionnaire data into a univariate outcome to represent a surrogate for the latent phenotype. However, when a causal variant is only associated with a subset of collapsed symptoms, the effect will be challenging to detect using the univariate outcome. We describe a more powerful strategy for genetic association testing in this situation that jointly analyzes the original multivariate symptom data collectively using a statistical framework that compares similarity in multivariate symptom-scale data from questionnaires to similarity in common genetic variants across a gene. We use simulated data to demonstrate this strategy provides substantially increased power over standard approaches that collapse questionnaire data into a single surrogate outcome. We also illustrate our approach using GWAS data from the Grady Trauma Project and identify genes associated with BDI not identified using standard univariate techniques. The approach is computationally efficient, scales to genome-wide studies, and is applicable to correlated symptom data of arbitrary dimension (thereby aligning with National Institute of Mental Health’s Research Domain Criteria).

## INTRODUCTION

Evidence indicates that common genetic variants should explain a sizeable role of the variation in many psychiatric disorders. For example, common variants are estimated to explain 40% of the heritability for bipolar disorder^1^, 21% of the heritability of depression^2^, 30-45% of the heritability of post-traumatic stress disorder (PTSD)^3-6^, and 50% of the heritability of autism spectrum disorder^7^. However, even in studies involving thousands of subjects, identification of specific common trait-influencing polymorphisms remains a challenge. To discover new associations, much attention has been spent on improving genotyping and sequencing technologies to interrogate more genetic variation; however, comparatively less attention has been afforded to thorough characterization of the underlying psychiatric phenotypes that are considered for genetic analysis.

In genetic analyses of a psychiatric phenotype, we often envision our outcome of interest as a single, measurable entity. In practice, we are rarely able to measure the outcome of interest directly and instead attempt to capture the true, latent phenotype via several connected but discrete measurements. As an example, psychiatric genetic studies attempt to account for the heterogeneity of symptoms found in a single psychiatric disorder by measuring the symptoms from several angles via a questionnaire or exam. For example, in studies of post-traumatic stress disorder (PTSD), researchers often measure the outcome using the PTSD Symptom Scale (PSS), which is a 17-item questionnaire for assessing and diagnosing PTSD according to the DSM-IV. Each item corresponds to a PTSD symptom and is rated from 0 to 3, with higher scores indicating greater symptom frequency/intensity ^8; 9^. Meanwhile, in studies of depression, many studies attempt to measure the phenotype using multiple symptom measurements from the Beck Depression Inventory-II (BDI). The BDI is a 21-item questionnaire with each question developed to correspond to DSM-IV diagnostic criteria for major depressive disorder. The answers to each question are scored from 0 to 3, with higher scores indicating more severe depressive symptoms ^10^.

Data captured by the PSS, BDI or other questionnaires can actually be considered a collection of interrelated multivariate phenotypes that, in the case of symptom scales, are usually ordinal in nature. The view of a mental disorder as a constellation of multiple correlated symptoms is aligned with the National Institute of Mental Health’s (NIMH) Research Domain Criteria (RDoC), which emphasize basic functional dimensions or mechanisms involved in psychopathology (e.g., fear, reward-seeking, attention, perception, arousal) rather than DSM or ICD diagnostic categories ^11^. Nevertheless, practical use of such multivariate symptom data in genetic analysis is complicated by the fact that standard statistical techniques for genetic analysis are generally univariate and designed to handle a single outcome at a time. To improve analytical utility, many questionnaires like the BDI and PSS were designed so that the multivariate symptoms are collapsed into a univariate phenotype for analysis. The simplest and most common collapsing method is unweighted summation of each question’s score ^10; 12-14^ into an univariate cumulative score. The cumulative score can then be treated either as a continuous outcome, or cutoffs can be applied to indicate presence/absence of disease symptoms.

An important issue with applying a univariate cumulative score in genetic analysis is that reducing multivariate information to univariate data nearly always comes at a cost. Carefully defining a phenotype is as vital in a GWAS as reliable genotyping; any association between gene and trait may be diluted by phenotypic heterogeneity. For example, if a gene were associated with a subset of the BDI questionnaire outcomes (e.g. a somatic symptom of depression-like changes in sleep patterns) but not other subsets (e.g. affective symptoms like mood or attitude), the magnitude of the overall effect size of the gene would be attenuated if the two subsets were combined into a univariate outcome measure.

A few key assumptions must be met in order for a univariate cumulative score to sufficiently summarize multivariate ordinal data. As noted by Van der Sluis et al.^15-17^, the three primary assumptions that must be met are: (1) the correlation between all questions in the questionnaire must be explained by a single (latent) phenotype; (2) the genetic effect must be on the latent phenotype; (3) the genetic effect—acting through the latent phenotype—must have identical effects on all of the questions in the questionnaire. For applied psychiatric phenotypes, it is more plausible that the assumptions are violated than maintained, a perspective that is supported by NIMH’s focus on RDoC. Depressive symptoms identified by the BDI might come from multiple sources (e.g. major depressive disorder, bereavement, post-traumatic stress disorder), violating the first assumption. The causal genetic effect might directly increase somatic symptoms of depression such as changes in appetite and sleep, but not impact mood, violating the second assumption. Alternatively, a variant might in fact affect each trait identified by every question, but have slightly different effect sizes on different questions. If any of these assumptions are not met, association analysis using the cumulative score will result in a substantial loss of power ^15; 17-19^.

A few alternatives have been presented to model the complex multivariate data captured within questionnaires. A popular type of approach is a data reduction method like principal component analysis (PCA), which relies on identifying a linear combination of the set of questionnaire responses that maximizes response variance across questions. Once the top few principal components are identified (i.e. those principal components that explain most of the questionnaire variance), association testing is performed between those top principal components and genotype ^20; 21^. However, PCA-based strategies that consider only high-variance principal components were recently shown to be generally suboptimal ^22^. As an alternative, Van der Sluis et al. ^17^ presented a multivariate gene-based association test by extended Simes procedure (MGAS) that combines the *P*-values obtained from standard, single-SNP association tests for each outcome to produce a single multivariate gene-based *P*-value. However, MGAS relies on permutations to establish significance, which make genome-wide analyses of psychiatric phenotypes cumbersome. Alternatively, Basu et al. ^23^ introduced a rapid multivariate multiple linear regression method (RMMLR), which operates on a MANOVA-based platform. However, while RMMLR establishes significance analytically, it cannot incorporate the important ordinal outcomes commonly measured in questionnaires and surveys.

To allow computationally-efficient and powerful genetic analysis of multivariate symptom data, we show in this paper that we can use a kernel distance-covariance (KDC) ^24-28^ method called the Gene Association with Multiple Traits (GAMuT) test ^29^, to assess association between high-dimensional symptom data and multiple variants (common or rare) in a gene. The framework is designed to test whether pairwise similarity in questionnaire responses is independent of pairwise genotypic similarity in a region of interest. The framework allows for an arbitrary number of categorical questions within the questionnaire as well as an arbitrary number of genotypes, thereby permitting gene-based or pathway-based testing of genetic variants. The method allows for covariate adjustment and is a closed-form test that yields analytic P-values, thus scaling easily to genome-wide analysis. GAMuT is therefore well-suited to facilitate research that is directly aligned with RDoC’s goal of encouraging investigation of biological, cognitive-behavioral, and self-report data using multivariate methods ^11^.

The remainder of this manuscript is organized as follows. We first provide a short overview of the GAMuT method and its features. We then present simulation work to demonstrate that the framework can be considerably more powerful than the standard univariate test based on a cumulative score derived from a questionnaire. We then illustrate the approach using a GWAS study of BDI scores collected as part of the Grady Trauma Project^30-32^. We finish with concluding remarks and discuss potential extensions to our approach.

## MATERIALS AND METHODS

### Overview of GAMuT

We provide a brief overview of the GAMuT method^29^ here and relegate the technical details of the procedure to the Supplementary Methods section. For a sample of *N* unrelated subjects, GAMuT examines the relationship between a set of *Q* questions (each question assumed to be an ordinal categorical variable with an arbitrary number of levels) and a set of *V* genetic variants within a gene or pathway of interest. GAMuT is motivated by the idea that, for a pair of individuals, increased genetic similarity at trait-influencing loci across a gene should lead to increased similarity of their questionnaire outcome data. Consequently, GAMuT constructs two different similarity matrices; one similarity matrix for the questionnaire outcomes and the other similarity matrix for the genetic variation within a gene. Each similarity matrix has *N* rows and *N* columns with individual elements of the matrix denoting the similarity (phenotypic or genetic) among different pairs of subjects. GAMuT creates a test statistic that evaluates whether the pairwise elements in the similarity matrix of questionnaire outcomes is independent of the pairwise elements in the genetic similarity matrix. The resulting test follows a known asymptotic distribution, which leads to easy and rapid calculation of p-values. GAMuT allows for questionnaire outcomes of arbitrary dimension and can further adjust for covariates.

### Simulations

We conducted simulations to verify that GAMuT properly preserves type I error (i.e., empirical size) and to assess power of GAMuT relative to standard association tests that treat questionnaire responses as a univariate outcome variable resulting from summing the responses into a continuous score. We briefly summarize the simulation design here and provide more comprehensive details in the Supplemental Methods section. We considered sample sizes of either 1000 or 2500 independent subjects. We performed simulations based on SNPs and LD patterns located within 2 kb up- and down-stream from *signal transducer and activator of transcription 3 (STAT3)*, a gene on chromosome 17q21.31 (see Supplementary Figure 1 for the MAF and pairwise LD structure of SNPs in *STAT3*). We generated simulated genotypes for all SNPs identified in HapMap within the *STAT3* gene (27 SNPs), but applied the testing approaches only to those SNPs that would be typed on standard genotyping arrays (14 SNPs).

We simulated multivariate questionnaire data to mimic the BDI questionnaire results obtained from Grady Trauma Project participants. The BDI consists of 21 groups of statements that reflect various symptoms and attitudes associated with depression. Each group includes 4 statements, which correspond to a scale of 0 to 3 in terms of intensity. The BDI is scored by summing the ratings given to each of the 21 items, yielding a cumulative score ranging from 0-63. To mimic BDI, we generated 21 ordinal responses using the observed distributions and correlations of these responses within the GTP BDI dataset. We show the correlation matrix among ordinal responses in Supplementary Figure 2 and the distribution of observations for each of the 21 ordinal responses in Supplementary Figure 3.

We applied GAMuT to 10,000 null simulated datasets to estimate empirical size. To investigate the performance of GAMuT under confounding and to assess whether the approach can successfully adjust for relevant covariates in this setting, we also tested empirical size by simulating questions under a confounding model where question responses were independent of genotype, but both questions and genotype were associated with a continuous covariate. For power models, we simulated data sets in which each of the 27 SNPs was modeled as being causal with effect size of the causal SNP on each question resulting in mean effect sizes with modest effect on the overall cumulative score. We varied the number of questions associated with the causal SNP, considering situations where 18/21, 12/21, and 6/21 questions were actually associated with the causal SNP.

Using the simulated data, we evaluated GAMuT using either projection matrices or linear kernels to model phenotypic similarity and using weighted linear kernels to model genotypic similarity (with weights based on sample MAF). We compared GAMuT to two standard approaches that use the univariate cumulative questionnaire score for inference: standard linear regression and kernel machine regression (KMR)^33^. Standard linear regression considers individual SNPs for analysis. KMR tests, on the other hand, jointly model multiple SNPs within a gene. KMR can be thought of as a specialized version of GAMuT that considers only 1 phenotype (the univariate cumulative sum of symptoms/questions) rather than the observed multivariate phenotypes. For KMR, we modeled genotypic similarity in in a fashion analogous to GAMuT by using a weighted linear kernel with weights based on sample MAF. Thus, comparison of GAMuT to KMR helps highlight the benefit of considering a multivariate questionnaire phenotype over a traditional cumulative-based score for gene-based analysis.

### Analysis of the Grady Trauma Project

Data used in our analyses were collected as part of the Grady Trauma Project (GTP), which investigates the role of genetic risk factors for psychiatric disorders such as PTSD and depression ^32; 34^. Participants in the GTP are served by the Grady Hospital in Atlanta, Georgia, and are predominantly urban, African American, and of low socioeconomic status. GTP staff approach subjects in the waiting rooms of Grady Primary Care, Obstetrics and Gynecology, and other clinics, obtaining their written consent to participate. In addition to collecting an Oragene salivary sample for DNA extraction, GTP staff conduct an extensive verbal interview, which includes demographic information, a history of stressful life events, and several psychological surveys, including the BDI.

The GTP initially genotyped participants on the Illumina HumanOmni1-Quad array to permit GWAS analyses. Applying standard GWAS quality control filters left 4,607 African-American subjects with good quality genotype data. Further removal of subjects who did not report at least one past trauma, subjects with missing BDI scores, or subjects with incomplete covariate data (age, gender, and the top ten principal components to account for ancestry) yielded a final sample size of 3,520 subjects.

For our sample, we used the support files provided by Illumina to identify 765,580 common genetic variants (MAF > 5%) that fell within 19,609 autosomal genes. We applied GAMuT to the BDI data using a linear kernel to measure pairwise phenotypic similarity in multivariate symptom scores. To measure genetic similarity, we used a linear genotype kernel within GAMuT and performed unweighted analyses as well as weighted analyses, with weights based on variants’ MAF or the variants’ estimated log odds ratios derived from external and independent GWAS studies of MDD, bipolar disorder, and schizophrenia that are available from the Psychiatric Genomics Consortium ^35-37^. For comparison, we also applied SNP-based linear regression and gene-based univariate KMR on the cumulative BDI score. For KMR, we applied the same genotype weighting schemes as used for the GAMuT analyses.

## RESULTS

### Type-I Error Simulations

Figure 1 shows the quantile-quantile (QQ) plots based on application of GAMuT, KMR, and linear regression to null datasets consisting of 1,000 or 2,500 subjects assayed for 21 BDI questions. For both sample sizes tested, GAMuT properly controls for type I error, even at the extreme tails of the test. KMR and linear regression, using the cumulative score approach, also demonstrated appropriate empirical size. Supplementary Figure 4 shows that residualization of questionnaire data prior to GAMUT analysis effectively controls for confounding that, unadjusted, would yield inflated results.

**Figure 1:**
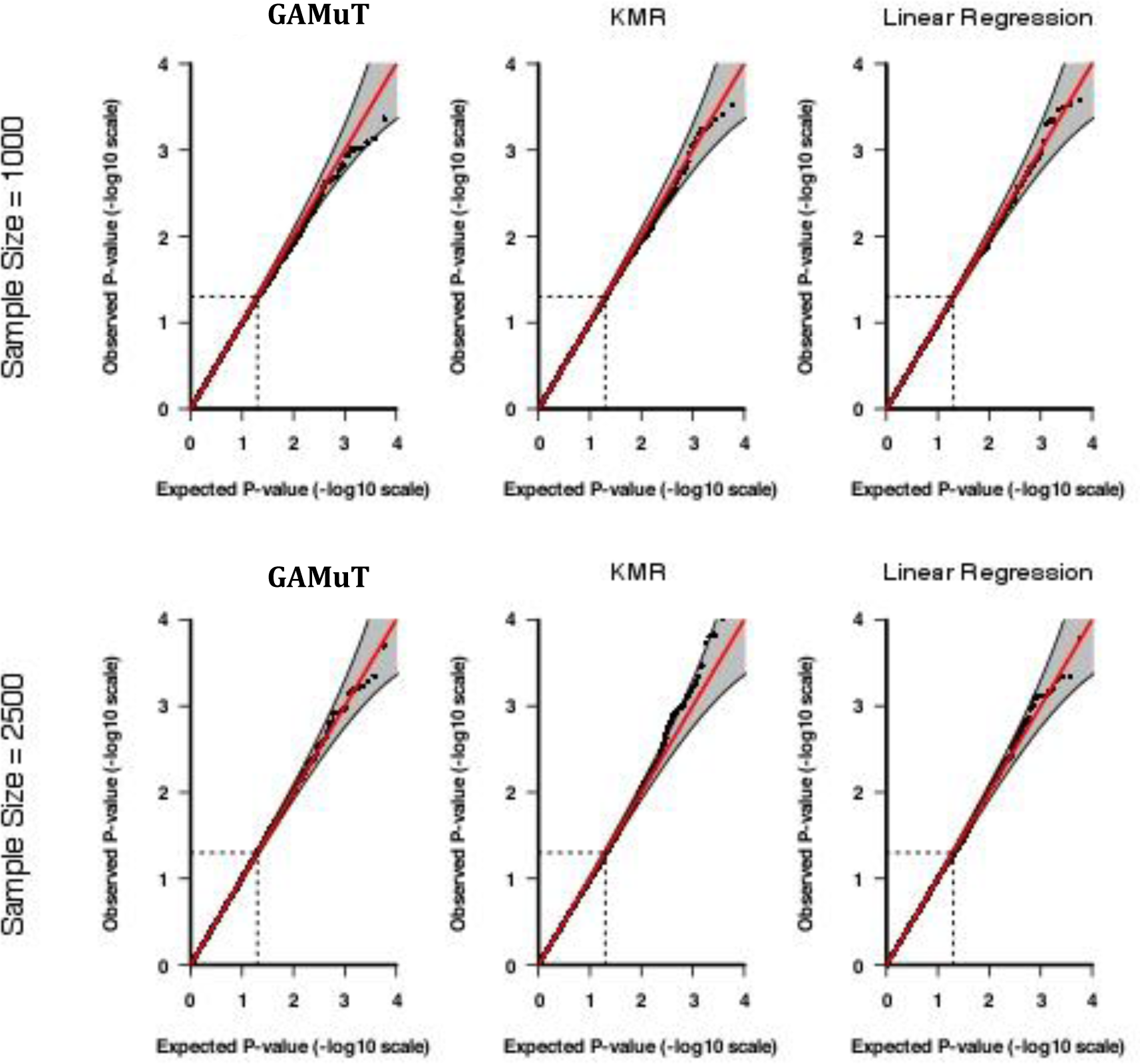
The QQ plots applying GAMuT, KMR, and linear regression to 10,000 simulated null data sets assuming a sample size of 1,000 (top row) and 2,500 (bottom row). For each simulation, 21 ordinal questionnaires were generated. For KMR and linear regression, the 21 questions were summed together to yield a single cumulative score.

### Power Simulations

Next we compared the power of GAMuT with univariate KMR and linear regression analyses in a series of simulation studies. For these power simulations, we set sample size to 1,000. Power was estimated as the proportion of *P*-values < 2.5x10^-6^ (reflecting a genome-wide correction for 20,000 genes) and was evaluated based on 500 replicates of the data per model. Figure 2 shows the power results. We plot power as a function of the causal SNP, where the causal SNPs are ordered by genomic location. The 14 genotyped SNPs (denoted by ‘x’ on the bottom of Supplementary Figure 1) were used to calculate test statistics, but all 27 SNPs were treated as causal in turn. Therefore, in situations where the causal SNP is not typed, we rely on correlation of the causal SNP with observed typed SNPs in STAT3 to gain statistical power. GAMuT offers considerably more power than the two competing univariate methods using cumulative scores for each of the three simulation models considered. When approximately half of the questions (12/21) are associated with the causal SNP, both KMR and linear regression observe nearly zero power to detect the effect; by comparison, GAMuT maintains power greater than 50% for 23 of the 27 causal SNPs. We observe a drop in power using GAMuT when nearly all of the questions (left column Figure 2) are associated with the causal variant compared with when a more modest number of questions are associated (middle column Figure 2). This pattern of decreased power when the proportion of associated phenotypes is close to 1 has been observed in other multivariate approaches, including multivariate analysis of variance (MANOVA) ^38; 39^. Regardless, our power results demonstrate the benefits of modeling the questionnaire data in a multivariate framework like that employed by GAMuT rather than using a traditional cumulative score.

**Figure 2:**
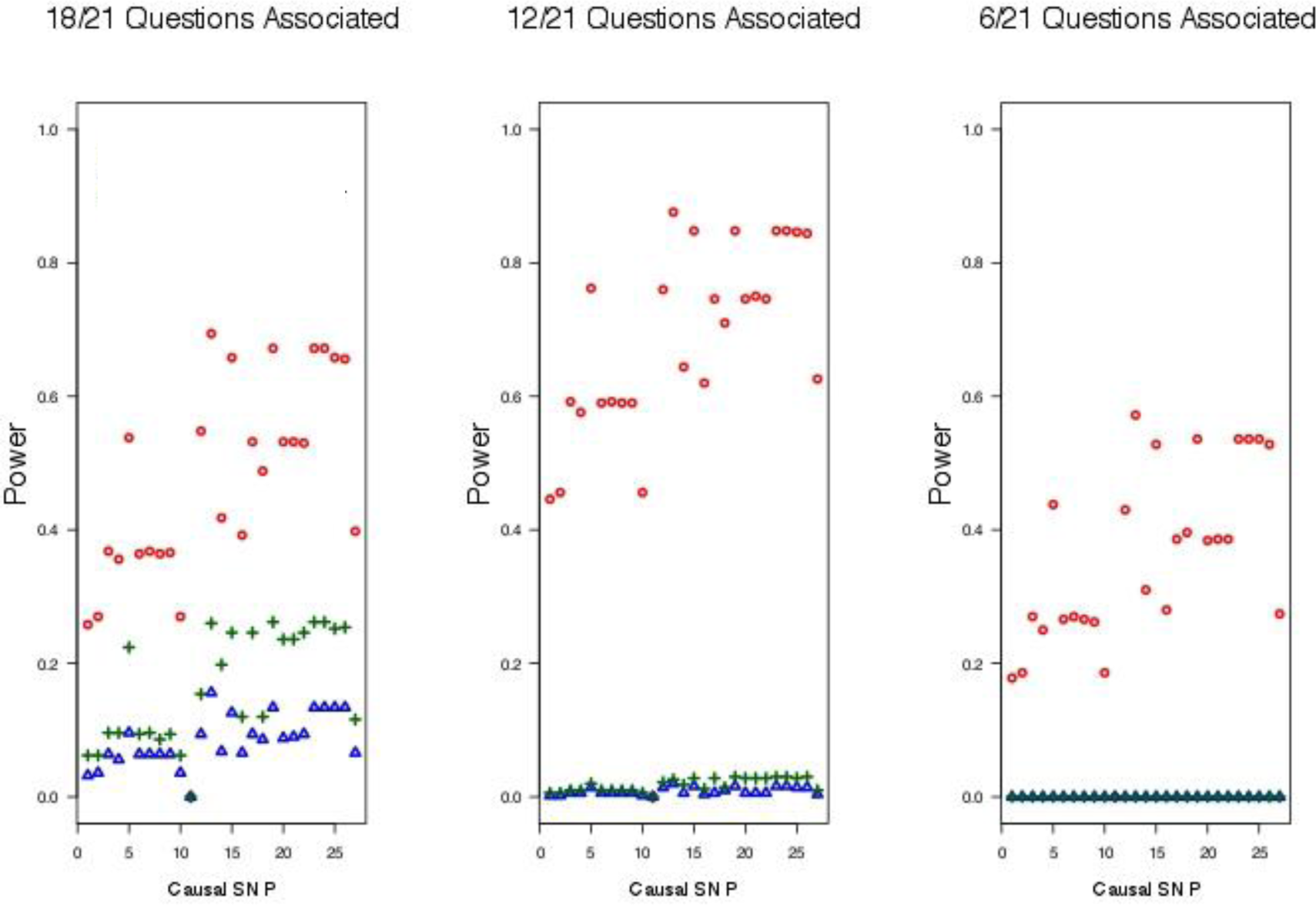
Power for GAMuT (red), KMR (blue), and linear regression (green) is plotted as a function of causal SNP. Left plot assumes the causal SNP is associated with 18 of the 21 BDI questions. Middle plot assumes 12 of 21 questions are associated with causal SNP. Right plot assumes only 6 of 21 questions are associated with the causal SNP. Sample size is 1,000.

### Application to Grady Trauma Project

We used the GTP dataset to test for associations between the BDI questionnaire and common variants in up to 19,609 genes. Prior to analyses, we controlled for gender, age, and ancestry in the 3,520 unrelated subjects. We applied GAMuT using a linear kernel to measure pairwise phenotypic similarity. We used several approaches for weighting SNPs, including MAF-based weights as well as external weights based on log odds ratio estimates from the PGC GWAS of MDD, bipolar disorder, and schizophrenia. For external weights, we note that not all GTP variants were present within the PGC GWAS results, and therefore the GAMuT analyses utilizing PGC-based genotype weights necessarily included fewer SNPs and corresponding genes than the analyses using MAF-based weights or no weights. Specifically, GAMuT analyses using PGC MDD weights involved 16,716 genes containing 469,582 SNPs. Meanwhile, GAMuT analyses using PGC bipolar disorder weights involved 16,761 genes containing 586,505 SNPs while analyses using PGC schizophrenia weights involved 18,067 genes containing 661,879 SNPs. For comparison with the GAMuT results, we ran univariate KMR using the cumulative BDI. For these KMR analyses, we employed the same genotype weighting schemes as used for GAMuT, and tested the exact same genes as tested in the GAMuT analyses. We also performed standard univariate linear regression of each of 775,255 common variants (SNP-level analyses) on the cumulative BDI score.

Since GAMuT and KMR analyze genes whereas linear regression analyzes SNPs, the multiple-testing adjusted significance thresholds differ between the former tests and the latter test. For each GAMuT and KMR analysis, we set a stringent study-wise significance threshold corresponding to a Bonferroni correction based on the number of genes tested (e.g., 0.05/19,609 = 2.55x10^-6^). Thus, the study-wise significance threshold differed depending on the particular genotype weights used, ranging from a threshold of 0.05/16,716 = 2.99x10^-6^ for PGC MDD weights to 0.05/19,609 = 2.55x10^-6^ for MAF-based weights and no weights. For all GAMuT and KMR analyses we considered *P*-values less than *P*<1x10^-4^ as suggestive. For SNP-based linear regression, we tested 775,255 SNPs across the genome. While we could apply the standard GWAS significance threshold of 5x10^-8^, we note that this threshold is more conservative than a Bonferroni correction based on the number of SNPs tested. Thus, for linear regression, we instead used a study-wise significance threshold of 0.05/775,255 = 6.45x10^-8^, and we considered *P*-values less than *P*<1x10^-6^ as suggestive.

We provide QQ and Manhattan plots for all GAMuT, KMR, and linear regression analyses of BDI in Supplementary Figure 5. We also list genes identified by GAMuT to be associated with BDI at study-wise or suggestive significance levels within Table 1. The GAMuT analyses of BDI identified one gene exceeding study-wise significance, while univariate KMR and linear regression of BDI did not detect any study-wise associated genes or SNPs. GAMUT found *ZHX2*, on chromosome 8, to be strongly associated with BDI (*P*=2.73x10^-6^), when using genotype weights based on estimated log odds ratios from the PGC GWAS for schizophrenia. We present QQ and Manhattan plots for this particular analysis in the first column of Figure 3. As noted in Table 1, *ZHX2* was also found to be highly suggestively associated with BDI when employing genotype weights based on the PGC GWAS of MDD (*P*=8.59 x 10^-6^). Previous research suggests a possible link between *ZHX2* and autism spectrum disorder ^40^. In comparison with the GAMuT analyses, KMR of cumulative BDI did not identify *ZHX2* as having even suggestive association (Table 1; Figure 3, middle column; Supplementary Figure 5b), and univariate linear regression revealed no SNPs suggestively associated with BDI within *ZHX2* or anywhere else across the genome (Table 1; Figure 3, last column; Supplementary Figure 5c).

**Table 1:** Full GAMuT Results for BDI (21 items)

| Gene | Chr | Number of variants | Genotype weights | GAMuT | KMR | Linear regression (minimum p-value of SNP in gene) |
| --- | --- | --- | --- | --- | --- | --- |
| ZHX2 | 8 | 97 | PGC SZ | $2.73 \times 10^{-6}$ | $4.42 \times 10^{-4}$ | $1.00 \times 10^{-3}$ |
| | | 76 | PGC MDD | $8.59 \times 10^{-6}$ | $1.36 \times 10^{-3}$ | |
| FAM43A | 3 | 115 | PGC BPD | $7.35 \times 10^{-5}$ | $4.15 \times 10^{-4}$ | $2.27 \times 10^{-5}$ |
| NUP214 | 9 | 19 | MAF-based | $7.97 \times 10^{-5}$ | $5.16 \times 10^{-3}$ | $5.57 \times 10^{-3}$ |
| E2F6 | 2 | 29 | PGC MDD | $9.45 \times 10^{-5}$ | $4.91 \times 10^{-2}$ | $3.32 \times 10^{-2}$ |
| GUK1 | 1 | 6 | PGC SZ | $9.54 \times 10^{-5}$ | $5.52 \times 10^{-3}$ | $2.34 \times 10^{-3}$ |
| SLC22A5 | 5 | 38 | PGC SZ | $9.69 \times 10^{-5}$ | $3.20 \times 10^{-3}$ | $3.98 \times 10^{-4}$ |

Genes with $P < 1 \times 10^{-4}$ identified in the GAMuT analyses are shown. GAMuT utilized a linear genotype kernel (possibly weighted) for all analyses. Of the genes listed, univariate KMR identified none using these same criteria, and standard linear regression identified no SNPs of suggestive significance (suggestive significance threshold for single SNPs: $P < 1 \times 10^{-6}$ ) within the gene. PGC MDD, PGC BPD, PGC SZ denote weights based on log odds ratios from the Psychiatric Genomics Consortium GWAS of major depressive disorder, bipolar disorder, and schizophrenia, respectively; MAF-based = weights based on minor allele frequencies of variants calculated using the Grady Trauma Project genotype data.

**Figure 3:**
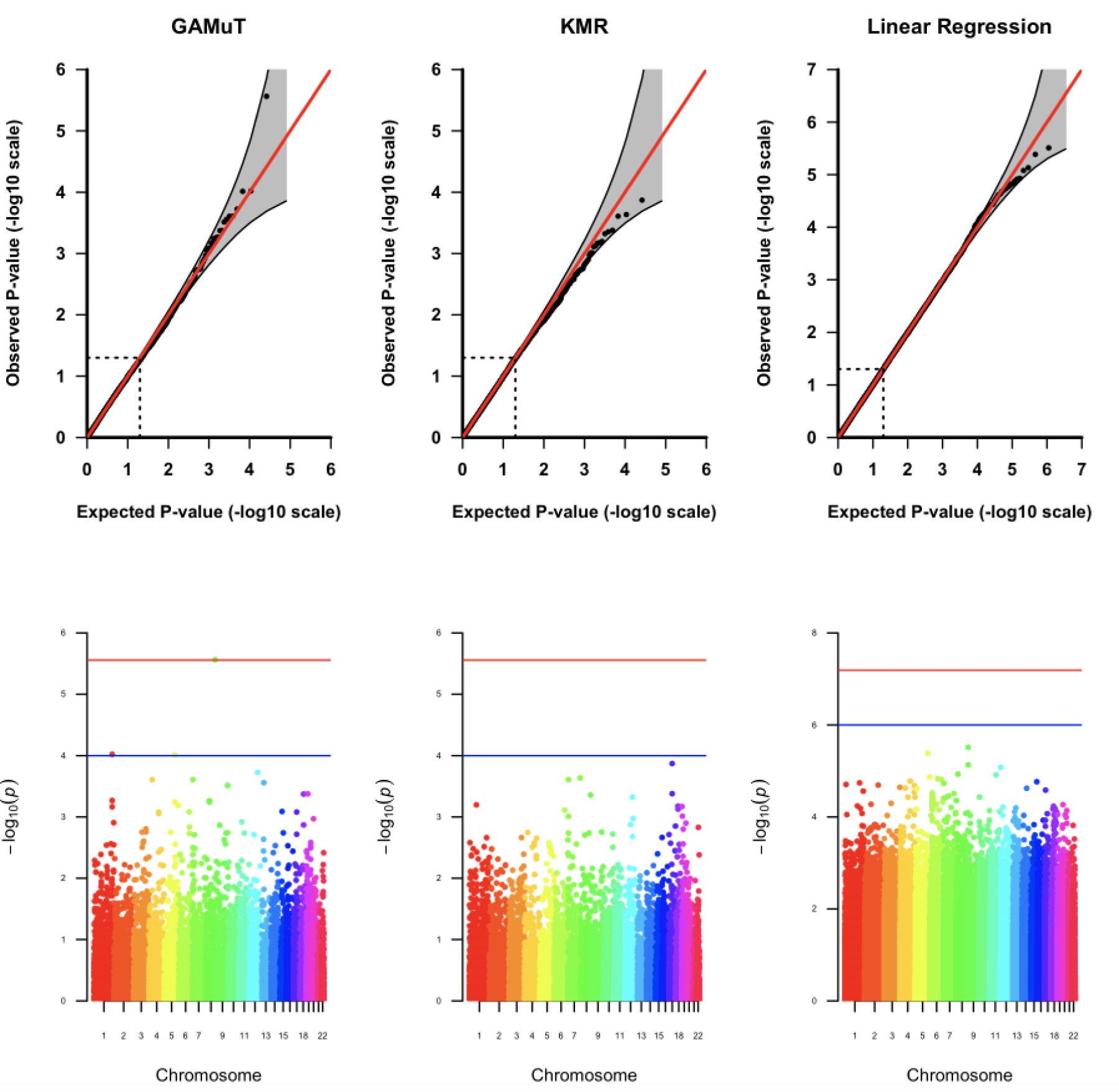
QQ and Manhattan plots for GAMuT, KMR, and linear regression analyses of BDI. The GAMuT analysis used a linear kernel to model phenotypic similarity and genotype weights derived from results of the PGC GWAS for schizophrenia. The KMR analysis also used weights based on the PGC GWAS for schizophrenia. In the Manhattan plots, the red line represents the study-wise significance threshold and the blue line represents the suggestive significance threshold. The study-wise significance thresholds for the GAMuT and KMR analyses are based on a Bonferroni correction for 18,067 genes tested, while the study-wise significance threshold for the linear regression analysis is based on a Bonferroni correction for 775,255 SNPs tested. In the Manhattan plot for the GAMuT results, the point exceeding the study-wise significance threshold is the -log_10_(*P*-value) for *ZHX2*, a gene on chromosome 8. These analyses used a sample of N = 3,520.

## DISCUSSION

As genetic studies of mental-health and psychiatric disorders increasingly shift to the study of high-dimensional symptom, questionnaire, and dimension data (such as those aligned with the NIMH RDoC), it is imperative to employ powerful statistical tests that maximize the possibility of novel genetic discoveries. Here, we have shown that multivariate methods like GAMuT are substantially more powerful for gene mapping of multivariate symptom data than standard methods that typically summarize such symptoms into a single univariate cumulative score for analysis. Methods like GAMuT that jointly model individual questionnaire outcomes are robust to phenotypic heterogeneity, in which a genetic risk factor only affects a subcategory within the questionnaire. In standard cumulative approaches, including KMR and linear regression, phenotypic heterogeneity can dilute the association between gene and trait, making the association extremely difficult to detect. While we focused here on gene-based studies of common variants, we note that our findings are generalizable to studies of rare genetic variation as well as studies of methylation patterns throughout the genome.

We applied GAMuT to the GTP dataset to test for associations between the BDI questionnaire and up to 19,609 genes. After controlling for important covariates, GAMuT found a strong association between BDI and *ZHX2* (*P*=2.73x10^-6^), which previous research suggests might be associated with autism spectrum disorder ^40^. In comparison, univariate KMR and linear regression did not identify *ZHX2* or SNPs within it to be associated with BDI, at even suggestive levels. This demonstrates through use of real-world data the capacity for multivariate methods like GAMuT to detect genotype-phenotype associations that would be missed using standard cumulative univariate approaches.

GAMuT derives analytic *P*-values based on Davies’ exact method, thereby improving computational efficiency and permitting application of the approach on a genome-wide scale. Like the popular KMR framework for univariate analysis, our approach allows for inclusion of prior information, such as biological plausibility of the SNPs under study. We provide R software implementing the approach on our website (see Web Resources). Computation run times for GAMuT are primarily dependent on sample size. Using a R script running single-threaded on a 1.7 GHz Intel Core i7 CPU processor, the time required for GAMuT to analyze 10 phenotypes for 1,000, 5,000, and 10,000 subjects is 0.52 seconds/gene, 13.2 seconds/gene, and 68.6 seconds/gene. Thus, genome-wide implementation is feasible particularly when high-performance cluster services are available.

## ACKNOWLEDGEMENTS

This work was supported by NIH grants GM117946, HG007508, MH071537, and AR060893. We thank Drs. David Cutler and Nigel Williams for their comments on a previous version of this manuscript. We appreciate the technical support of all of the staff and volunteers of the Grady Trauma Project. Most importantly, we are extremely indebted to and appreciative of the time and effort given from all of the participants of the Grady Trauma Project.

## WEB RESOURCES

Epstein Software: https://github.com/epstein-software

OMIM: http://www.omim.org

